# The war of corals: patterns, drivers, and implications of changing coral competitive performances across reef environments

**DOI:** 10.1101/2021.12.06.471466

**Authors:** Mohsen Kayal, Mehdi Adjeroud

**Affiliations:** ENTROPIE, IRD, CNRS, IFREMER, Université de la Nouvelle-Calédonie, Université de la Réunion, Noumea, New Caledonia; Laboratoire d’Excellence “CORAIL”, Paris, France; ENTROPIE, IRD, CNRS, IFREMER, Université de la Nouvelle-Calédonie, Université de la Réunion, Perpignan, France; PSL Université Paris, USR 3278 CRIOBE - EPHE-UPVD-CNRS, Perpignan, France

**Keywords:** Coral reef, Competition, Territorial war, Niche segregation, Species coexistence, Global change

## Abstract

Amidst global environmental changes, predicting species responses to future environments is a critical challenge for preserving biodiversity and associated human benefits. We explored the original idea that coral competitive performances, the ability of corals to preempt ecological space on the reef through territorial warfare, serve as indicators of species’ ecological niches and environmental windows, and therefore, responses to future environments. Our surveys indicated that coral performances varied with taxonomic-identity, size, and position along environmental gradients, highlighting complex interplays between life-history, warfare-strategy, and niche segregation. Our results forewarn that growing alterations of coastal environments may trigger shifts in coral dominance, with decline of major reef-building taxa like acroporids, and underscore the importance of restraining human impacts for coastal resilience. Our empirical approach untangles the complexity of species’ battle-like interactions and can help identify winners and losers in various communities caught in the interplay between ecological niches, environmental windows, and global changes.

## Introduction

Predicting how global environmental changes will affect species performances in the future is crucial to anticipating biodiversity declines and defining sustainable management. However, large uncertainties blur current predictions of ecosystem trajectory in future environments [1,2]. Finding effective metrics of species responses to changing environments is key, particularly for vulnerable ecosystems in need of rapid intervention, and in developing and island nations where high reliance on natural resources exacerbates socio-ecosystem vulnerability [3–5]. This is particularly true with coral reefs, which support prolific marine life and coastal livelihoods yet stand at the frontline of declining ecosystems due to rapidly altering coastal environments [1,6–9]. Reef degradation from growing coastal development, pollution, fishing, and climate change predominantly involves a gradual decline in coral abundance, composition, and size, with the progressive loss of vulnerable species, particularly at sensitive life-stages, altering key ecosystem functions [7,10–13]. Recent demographic modeling approaches allow characterizing these dynamics on reefs with fine-scale monitoring data [2,14– 16]. However, only a few eminent sites, representing an infinitesimal proportion of reefs, benefit from the necessary level of scientific knowledge, leaving out most coral reef ecosystems from such quantitative diagnosis.

As an alternative to using demographic modeling, we hypothesize that coral competitive performances, the ability of corals to preempt space on the reef substrate through territorial warfare, could be used as a proxy of species ecological success in different environments. Competition for space and other limiting vital resources is a key process shaping ecological communities in coral reefs, notorious for biodiversity and biotic interactions, and where competition warfare can drive community shifts and ecosystem collapse in altered environments [17–19]. We stipulate that differences in competitive performances across environments can provide insights on species ecological niches, optimal environmental windows and, therefore, potential response to future conditions.

On reefs, direct competition for space is ubiquitous where neighboring organisms grow into physical contact, species inevitably engaging in warfare for survival and ecological dominance. On the front line (a.k.a. the battle-zone), corals have the capacity to invade opponent territories by killing enemy living tissues in their vicinity. This battle predominantly takes two forms with either smothering by growing over (a.k.a. overgrowth) or disintegrating using specialized nematocyst-rich attack tentacles (a.k.a. overreach), or a combination of the two (figure 1). In theory, the magnitude of killing varies with fixed characteristics of species life history such as attack mechanism, strength, and reach [14,20–24], but also with additional processes that vary in time, space, and across life-stages such as growth rates and metabolic states of corals [25–29]. As such, while a clear hierarchy of competitive dominance among coral species can be established in a given environment, competitive outcomes in fact appear spatio-temporally dynamic [30–34], and are therefore expected to differ in the future with changing reef environments. Because corals are slow growing, habitat-forming species at the foundation of reef ecosystems, even small differences in species abilities to preempt reef space can have strong implications for reef ecosystem structure and functions, and associated services to society.

**Figure 1.**
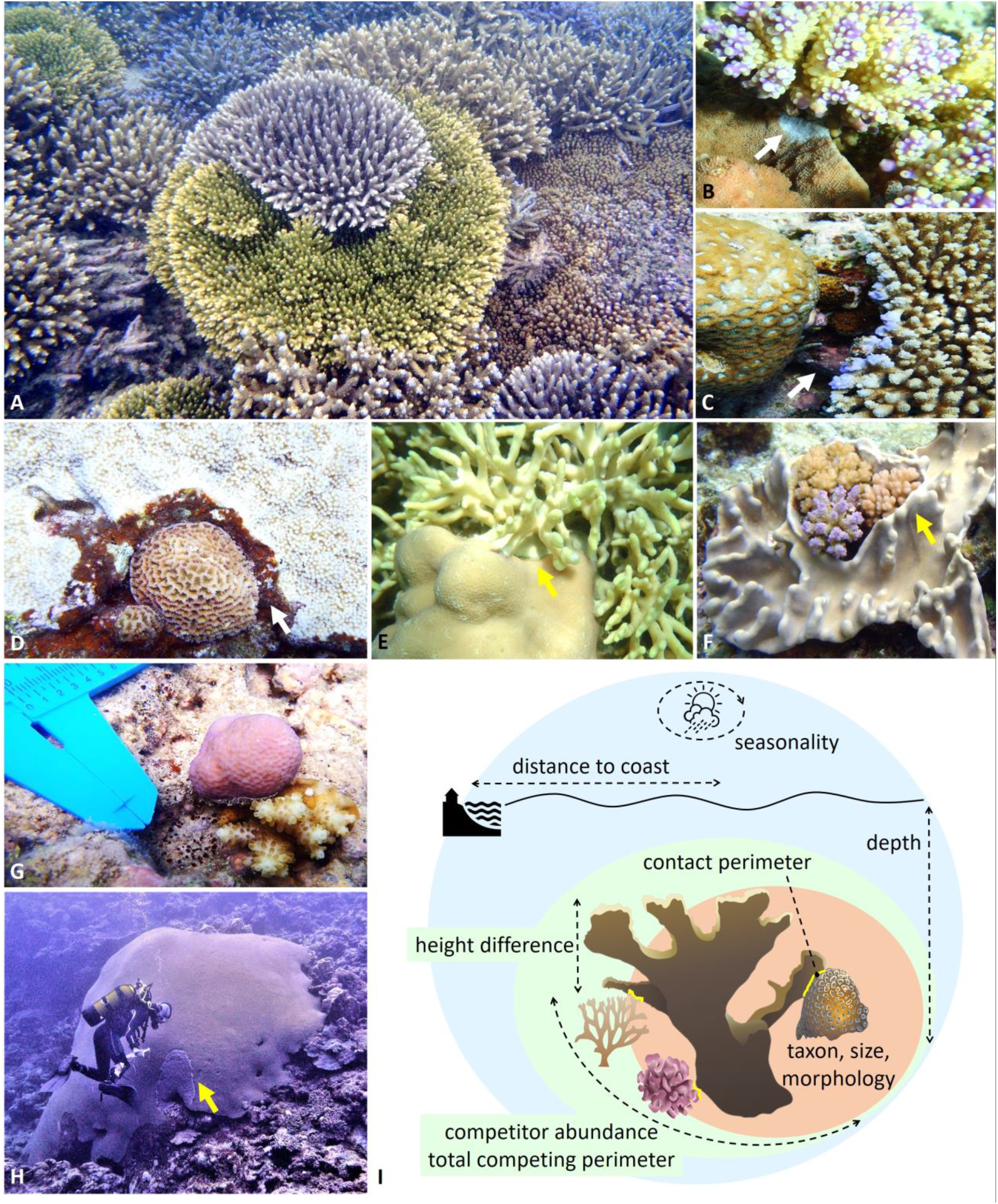
Photographs illustrating encountered coral competitive interactions (A-H), and schematic (I) indicating how they were characterized by taking into account a set of factors relating to individual organisms (beige), interactions (green), and environments (blue). By quantifying the dead-zones left by sweeper attack-tentacles on opponent coral skeletons in the aftermath of competitive battles, overreach distances (white arrows) reflect short-term competitive outcomes resulting from recently deployed assaults (hours to months preceding observations). In contrast, given the slow growth of corals, overgrowth distances (yellow arrows) often integrate competitive interactions over several years. See appendix 1 for further information, and figure S2 for visualizations of the raw data.

We used observations of coral competitive interactions across the south-western reef system of New Caledonia as indicators of the capacity of coral species to prevail in different environments. The island nation is surrounded by large extents of biodiverse coral reefs, characterized by diverse habitats distributed along pronounced coast-to-ocean gradients associated with natural environmental variability amplified by human impacts [35,36] (figure S1). The near shore is most exposed to terrigenous inputs of freshwater, nutrient, and sediment, as well as anthropogenic impacts such as pollution and fishing, and undergoes higher environmental variation with seasons and weather conditions [37–40]. Moving towards the ocean, marked increases in water quality and diminishing human pressure is observed. Portions of the reefs are classified as UNESCO World Heritage site due to their outstanding character for global coral reef conservation. Zonation in the relative abundance of coral species along the coast-to-ocean gradient indicates some degree of niche segregation among dominant taxa [36]. However, the ecological mechanisms underlying such spatial patterns largely remain to be comprehended. Because coral species are expected to occupy delimited ecological niches distinguished by environmental preferences [7,29,41], we tested whether the spatial distribution of ecological windows would be reflected in coral competitive outcomes. In general, a deeper understanding of coral competition can help a better characterization of coral life-strategies, which at this stage predominantly relies on qualitative assumptions of species competitive abilities based on taxonomic traits and demographic performances [28,42]. A better understanding of coral competition can also inform on the ecological processes underlying species coexistence and the exceptional biodiversity of coral reefs. We illuminate our findings with analogies to warfare theory and lessons from human history to untangle the complexity of coral competitive interactions, and discuss implications for coral performances in changing reef environments.

## Methods

We evaluated coral competitive performances by inspecting natural occurrences of direct physical interactions among corals, as well as with other sessile benthic species (figure S2). Surveys were performed on 20 sites distributed along pronounced cross-shelf environmental gradients (figure S1), and over a five-month period to capture seasonal variability with a shift from the warm (January) to the cold (June) season (decreasing water temperature, 27-23°C). For each interaction haphazardly encountered, the taxonomic identities, morphotypes, and three-dimensional size (length, width, and height) of each organism (the focal coral plus all its direct competitors) were recorded, along with the contact perimeter characterizing the battle zone, and overreach and overgrowth distances as short- and long-term metrics of competitive outcomes (figure 1). Because coral demographic performances in survival, growth, and reproduction vary with size [28,43], changes in coral abilities to defend their territories were related to species fitness (i.e. chances of ecological success). We focused specifically on direct competition for space (a.k.a. territorial war), leaving aside indirect competition for light, food, and other resources.

As competitive interactions regularly consisted of bilateral attacks, net overreach and overgrowth performances were calculated as the difference between the observed maximum conquered and ceded distances along the frontline. These measurements of net performances differ from most studies on competition, which typically characterize species interactions as a simplistic binomial (win or loss) or trinomial (win or loss or standoff) outcome (e.g. [23,24,31– 33]). Further characteristics of the interaction and environment that could also potentially influence competitive outcomes, such as distance to the coastal city Noumea, water depth, competitor abundance, total competing perimeter, and differences in height among competitors, were also recorded (figure 1I, table S1).

Because coral performances were expected to be influenced by various ecological factors acting at different scales and that in concert shape species responses (figure 1I), we used generalized additive models to characterize changes in competitive outcome as a function of different candidate ecological descriptors (table S1) in a non-linear, multi-dimensional account [44]. Model parametrization was designed to capture a set of common ecological processes regulating coral performances, such as size- and density-dependence (e.g. covariates *Coral-size* and *Competitor-abundance*), as well as taxonomic deviations in such processes (e.g. interaction *Coral-size* × *Coral-taxon*) to account for evolutionary differences among species [28]. The degree of non-linearity of model terms was optimized based on semi-parametric spline-penalization (see [28,45] for details), and non-significant model terms were sequentially excluded during the model selection process [44] (figures S3 and S4). Among the multitude of possible models resulting from combinations of the explanatory variables, the models best describing competitive outcomes were identified using Akaike information criteria, a measure of trade-off between model performance and complexity [46]. The final models (tables S2 and S3) explained variability in coral competitive performances at 66.6% in terms of net overreach and 79.0% in terms of net overgrowth (figures S3 and S4). This is relatively high compared to previous attempts (e.g. [23,24,32]) and considering the many additional biological and environmental factors that may influence competitive outcome between two living organisms (genotype, age, health, metabolic state, disturbance history, etc.), suggesting that our models accounted for key ecological gradients influencing coral competitive performances in our study system. Restricting data to the most abundant taxa resulted in similar patterns (figures S5 and S6), confirming the prevalence of the identified mechanisms among dominant species.

A total of 1073 competitive interactions were recorded encompassing 41 taxa and 8 morphotypes (table S1). All surveys were performed by the same observer using SCUBA, occasionally assisted by another diver. Analyses and graphing were coded in R statistical software complemented by the mgcv package [44]. Over-dispersed variables were log-transformed, and model residuals were systematically checked for normality and homoscedasticity.

## Results and Discussion

### Identifying drivers of coral competitive performance

Of the 1073 coral competitive interactions inspected, 84.3% (905) involved traces of tentacle deployment along the frontline, out of which 7.4% (67, or 6.2% of all interactions) were bilateral. In 8.9% (6) of these bilateral overreach attacks, net space intrusions were tied between competitors (i.e. net overreach = 0). Similarly, 69.8% (749) of all interactions involved overgrowth, out of which 2.0% (15, or 1.4% of all interactions) were bilateral. In 13.3% (2) of these bilateral overgrowth attacks, net space invasions were tied between competitors (i.e. net overgrowth = 0). Only 0.5% (5) of all interactions were characterized as standoffs for both overgrowth and overreach, demonstrating the complementary nature of these two metrics to assess competitive wars among corals.

Coral competitive performances varied with attributes relative to individuals, interactions, and environments, highlighting the interactive importance of biological and environmental factors in defining competitive outcomes. Overgrowth and overreach performances were both contingent on taxonomic identity, morphology, size, competitor abundance, and shelf-position, whereas perimeter of contact and height differences only influenced overgrowth, and day of year only overreach (tables S2 and S3). As a metric of short-term competitive interactions, overreach reflects recently deployed battle strategies such as spontaneous attack-tentacle developments into opponent territories, brief skirmishes along the frontline that were seasonally variable in several taxa (figure 2C) and may reveal transitory in long-running competitive battles [22,25,30,32,34]. In contrast, as a more integrated measure of competitive interactions over time, overgrowth accounts for additional ecological mechanisms that prevail across successive battles in war strategy. This includes the capacity to sustain siege and lead large battlefields, sometimes for long times and simultaneously on multiple fronts (figure 1), performances that rely heavily on resource provisions and differed across taxa.

**Figure 2.**
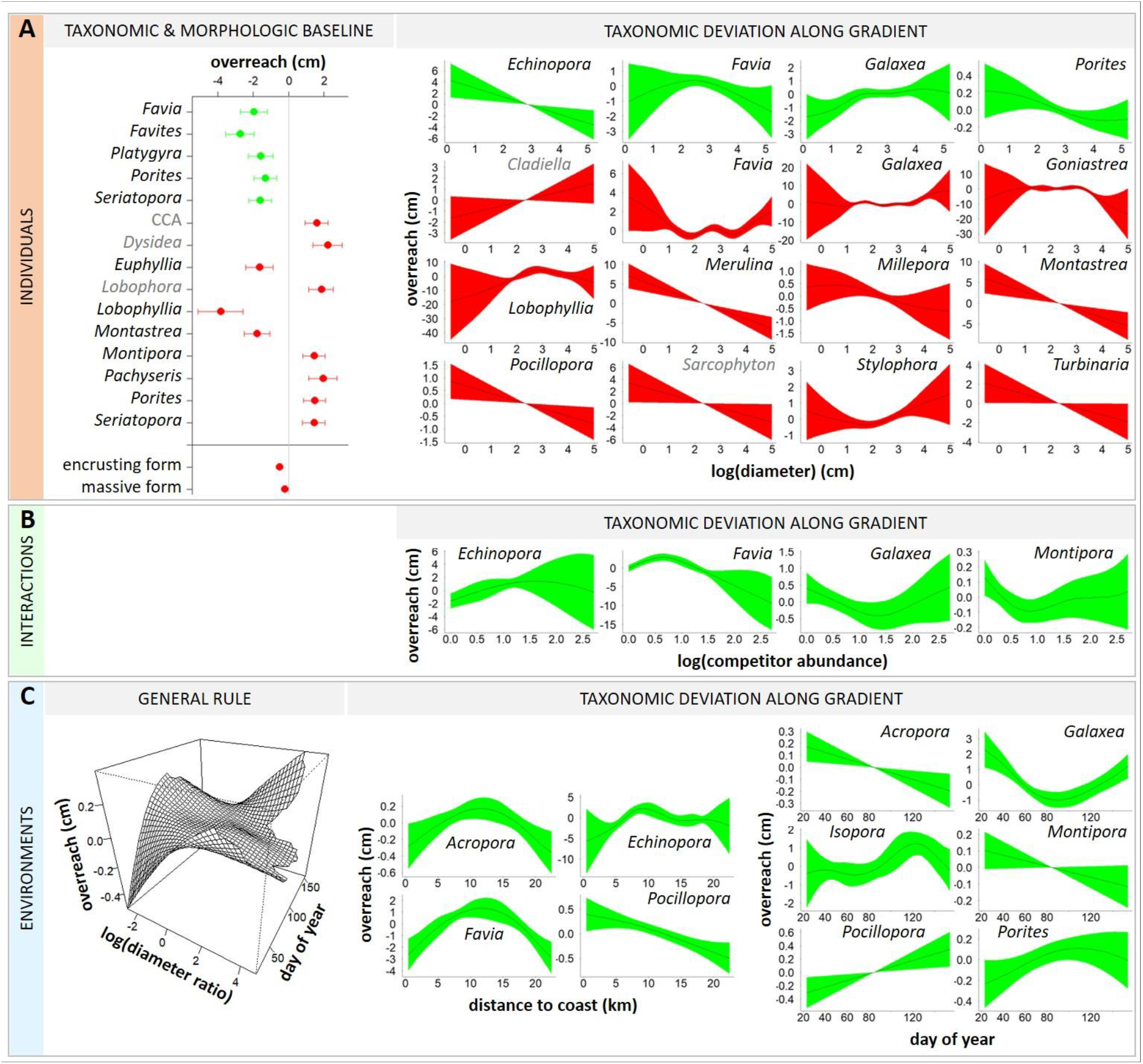
Changes in coral competitive performance as measured by net overreach distance along multiple ecological gradients. Plots illustrate partial contributions of different covariables to variation in net overreach of focal corals (mean ± standard error). Covariables are organized by scale, characterizing which organisms are involved (A, individuals), and how (B, interactions) and where/when (C, environments) the interactions occur. Covariables measured on focal corals are displayed in green (e.g. a positive effect of focal coral diameter on focal coral performance) and those on competing organisms in red (e.g. a negative effect of competitor diameter on focal coral performance). Taxonomic and morphologic baselines identify differences in performance among species and growth-forms once the effects of other ecological gradients are accounted for. Three-dimensional plot illustrates the interactive effects of two ecological gradients on the response of all species, while other plots indicate deviations specific to some taxa and growth forms. Note differences in axes ranges. Texts in grey distinguish non hard-coral species (CCA for crustose coralline algae). Only significant effects are illustrated (table S2).

### Individual-level attributes

Coral overgrowth and overreach performances differed among taxa as expected for species exhibiting contrasting life history traits, with differences in growth form and rate, in tentacle size and reach, etc. [23,34,42]. Yet, contrasting responses to ecological gradients provided deeper insights into distinct life-strategies as reflected by different patterns of competitive performance across life-stages, as well as contrasting susceptibilities to environmental variation as reflected by segregated environmental optima. Larger corals generally exhibited higher net overgrowth, though many taxa deviated from a common size-dependent pattern in overgrowth and overreach (figures 2A and 3A). While evidence of size-dependent variability in coral competitive performance is not new [25,26], our comparative study indicates maximum competitive capacities occur at different stages among species, suggesting differences in size-specific investments in competition. Some taxa showed higher overgrowth and overreach at small sizes, potentially in a strategy to secure enough space early on, until reaching a size-refuge that guarantees survival and investment in other demographic processes such as reproduction [28,33,43]. This was the case for *Porites*, in which competitive performances declined with colony size (−0.3 cm in net overreach and -1.5 cm in net overgrowth across the size-range), with an inflection point at a size of ∼15 cm diameter (figures 2A and 3A). Other taxa performed better at intermediate or larger sizes, which corresponds with higher ability in allocating large energetic resources to competitive battles. A marked positive effect of colony size was detected in mean overreach of *Merulina* (+14 cm across the size-range), *Montastrea* (+12 cm), and the soft-coral *Sarcophyton* (+7 cm), as well as in mean overgrowth of *Goniastrea* (+11 cm), *Hydnophora* (+9 cm), *Merulina* (+10 cm), and the soft-coral *Nephthea* (+70 cm). The latter taxon reveals being a particularly fierce competitor of reef-building corals with an unmatched ability to overgrow them (figure 3A) [21].

**Figure 3.**
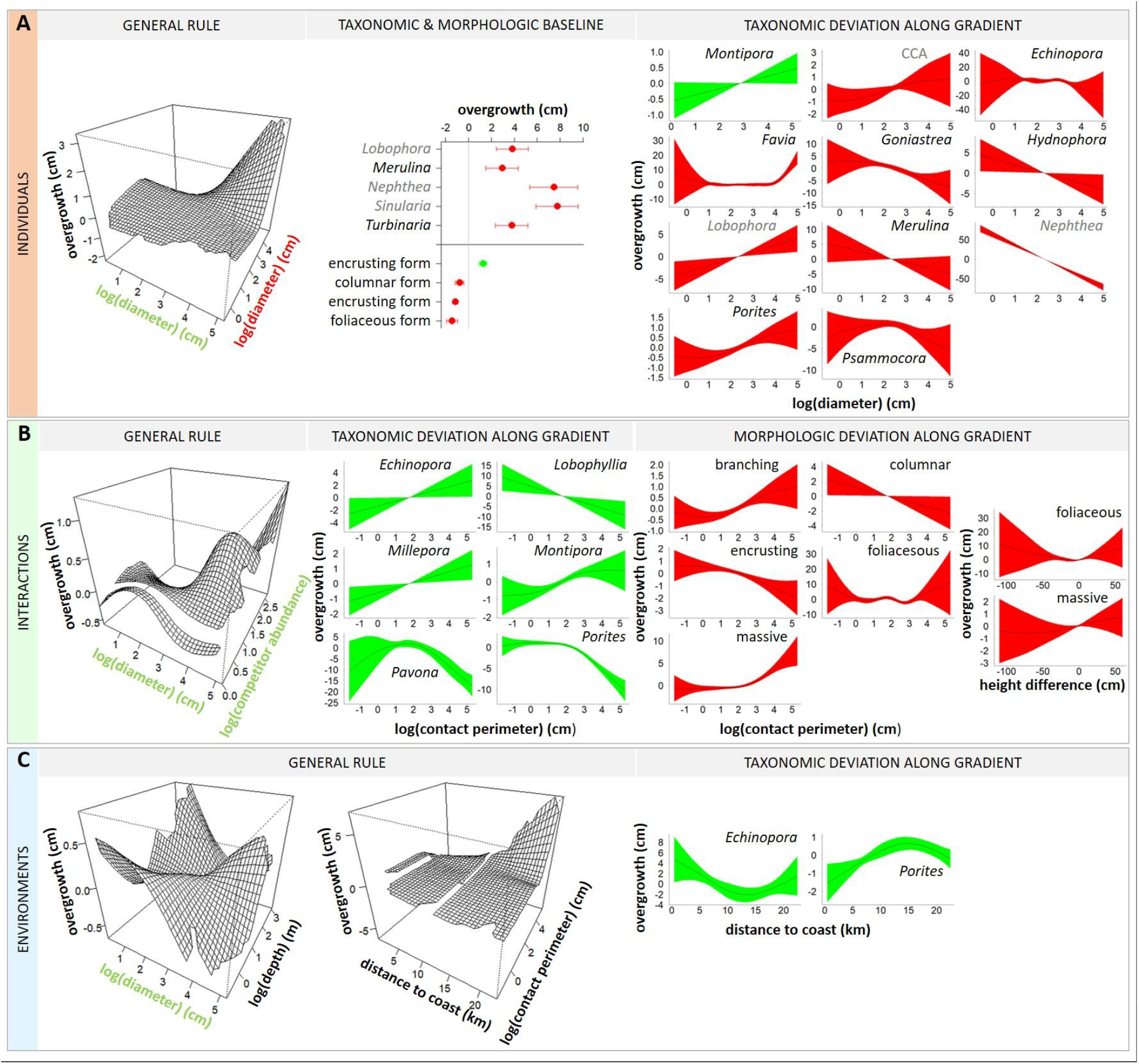
Changes in coral competitive performance as measured by net overgrowth distance along multiple ecological gradients. Plots illustrate partial contributions of different covariables to variation in net overgrowth of focal corals (mean ± standard error). Covariables are organized by scale, characterizing which organisms are involved (A, individuals), and how (B, interactions) and where/when (C, environments) the interactions occur. Covariables measured on focal corals are displayed in green (e.g. a positive effect of focal coral diameter on focal coral performance) and those on competing organisms in red (e.g. a negative effect of competitor diameter on focal coral performance). Taxonomic and morphologic baselines identify differences in performance among species and growth-forms once the effects of other ecological gradients are accounted for. Three-dimensional plots illustrate the interactive effects of two ecological gradients on the response of all species, while other plots indicate deviations specific to some taxa and growth forms. Note differences in axes ranges. Texts in grey distinguish non hard-coral species (CCA for crustose coralline algae). Only significant effects are illustrated (table S3).

### Interaction-level attributes

Coral competitive outcomes were influenced by several characteristics of species interactions, namely competitor abundance (a.k.a. number of enemies), contact perimeter (a.k.a. battlefield stretch), and height difference among competitors (a.k.a. unequal battlegrounds). Competitor abundance was associated with changes in overreach performances of four coral taxa (figure 2B), and generally influenced size-specific overgrowth, with larger corals showing higher capacities in leading multi-front wars (figure 3B). Larger battlefields were associated with higher overgrowth in some taxa, such as *Millepora* and *Montipora* (+2 cm in net overgrowth), showing high capacities in waging large-scale competitive endeavors to the detriment of others such as *Porites*, in which a threshold in the capacity to hold space against competitors was observed (−10 cm in net overgrowth, with again a size-threshold at ∼15 cm diameter, figure 3B). Overall, extended battlefield perimeters were associated with higher overgrowth performance in encrusting and columnar species and decreasing in branching and massive taxa, reflecting evolutionary differences in competitive abilities among morphological groups [24,42].

### Environmental attributes

Battlefield environments also influenced coral competitive performances as reflected by the effects of shelf position, depth, and time, revealing differing environmental preferences among taxa (tables S2 and S3). Cross-shelf variability in coral performances was detected in five taxa, among which peak overgrowth and/or overreach performances were spatially segregated. *Pocillopora* showed higher net overreach (+1 cm) near the coast, whereas *Acropora* (+0.6 cm) and *Favia* (+4 cm) peaked in mid-lagoon, *Porites* exhibited highest overgrowth (+2.0 cm) towards the barrier-reef, and *Echinopora* showed contrasting spatial patterns between overgrowth and overreach metrics (figures 2C and 3C). Similarly, overreach performances varied seasonally in six taxa with a marked temporal segregation (figure 2C). *Acropora* (+0.4 cm) and *Montipora* (+0.2 cm) showed higher performance in warm season, *Pocillopora* (+0.7 cm) in cool season, *Isopora* (+1.5 cm) and *Porites* (+0.3 cm) during the inter-season, whereas *Galaxea* (+3.5 cm) peaked in each season. While the spatial differences in competitive performances among taxa confirm contrasting environmental optima along the coast-to-ocean gradient, the temporal patterns identified may reflect differences in environmental preferences *per se* (e.g. differing temperature optima) or different timings of investments in other demographic processes such as growth and reproduction (i.e. differing temporal windows). Indeed, territorial wars as well as growth and reproduction are energetically costly processes, and some corals may show temporal tradeoffs in their investment in these endeavors [22,26,28,32]. For example, in New Caledonia as in the neighboring Great Barrier Reef, acroporids including *Acropora* and *Montipora* reproduce at the onset of the warm season following a 6-month period of gametogenesis, whereas other species such as *Pocillopora* reproduce throughout the year [47– 50]. In addition, portions of the temporal trends identified may be attributable to response to external stimuli, which may explain the higher overreach observed in *Porites* during the inter-season between the performance peaks of acroporids and *Pocillopora*, and conversely high overreach in *Galaxea* during both seasons potentially as retaliation to attacks from these dominant taxa [32,36].

### Species baseline attributes

When the effects of ecological gradients were isolated, species with encrusting and massive morphologies were associated with higher overreach (figure 2A), whereas encrusting, columnar, and foliaceous species exhibited higher overgrowth (figure 3A). These differences may reflect differing evolutionary pathways among species. In contrast to other growth forms that enable refuging via vertical ascension, encrusting species are fully exposed to competition and their survival relies fundamentally on their capacity to preempt space on a two-dimensional substrate [14,22,33]. Similarly, several coral species with massive growth forms exhibit large polyps able to rebuff competitors on longer distances by developing long-range tentacles (figure 1) [20,23,24,34]. A maximum overreach distance of 5.7 cm performed by a massive *Galaxea* on a branching *Pocillopora* was recorded in this study, and the three massive taxa *Euphyllia, Lobophyllia*, and *Montastrea* exhibited high baseline overreach (figure 2A). Noticeably, marked negative effects of competitors on coral overreach and overgrowth were only detected from hard-and soft-corals, whereas interactions with ascidians, sponges, and algae were characterized solely by positive deviations from the average patterns (figures 2A and 3A). This suggests a less substantial effect of chemical wars alone as employed by these later taxa, compared to additional uses of physical wars involving tentacle-attacks as deployed by cnidarians, in direct competitive interactions [27,51,52].

### Comprehending coral competitive interactions

Our study shows that coral competitive performances are governed by a complex interplay between who is involved and how, where, and when the interactions occur, with outcomes in terms of net overreach and overgrowth that are largely predictable (figures 2 and 3). Species baseline performances associated with inherited taxonomic traits (e.g. tentacle reach) are modulated by a set of ecological gradients related to intrinsic characteristics of organisms (e.g. size, evolutionary life-strategy) as well as extrinsic features of their interactions and environments that vary in time and space (e.g. competitor abundance, contact perimeter, seasonality). The mechanisms governing coral competitive performances are therefore fundamentally analogous to those prevailing in human warfare where concerted military power (weapon abundance, deadliness, and reach), war strategy (attack, defense, skirmish tactics), battle characteristics (stretch of battlefronts, duration of conflicts, number of enemies and allies), and battlefield features (battleground evenness, weather conditions) influence outcomes [53,54]. Notorious instances when battle characteristics and environmental condition influenced war outcome and sealed the fate of human history include the deleterious effects of multi-front wars for Napoleon’s endeavor to expand the French empire across Europe between 1805 and 1815 [55], and the contribution of wintery weather conditions to Hitler’s army’s defeat at the Soviet frontline in 1941 [56]. Another historical example relates to the Battle of Agincourt in 1415, a turn in the Hundred Years’ War for the dominion of France and England, where muddy terrain following rainfall severely handicapped the heavily armored knights of the numerically superior French army to the advantage of Henri V [57].

### Niche segregations and implications in changing environments

The contrasted responses of coral taxa as identified across multiple ecological gradients provide new insights into the variety of mechanisms underlying niche segregation in biodiverse species assemblages. Indeed, the diversity of ecological windows occupied by species is reflected in the contrasting environmental preferences revealed by differing performances in time and space, as well as the divergent evolutionary pathways as indicated by different responses to individual and interaction level attributes (figure 4). These differences may explain how the species coexist as a result of distinct demographic life-strategies, environmental heterogeneity, and competitive interactions, resulting in the exceptional biodiversity observed on reefs [17,23,28,42]. Nevertheless, several key coral taxa were sensitive to environmental variability as reflected by distance to the coast and seasonality, indicating that their competitive success, and perhaps overall fitness, may be affected by alterations of coastal environments. Our findings suggest that warmer oceanic conditions, similar to those presently observed in summer, may advantage higher competitive performances of acroporids to the detriment of pocilloporids, although further anthropization of coastal habitats, currently restricted to the coastline, benefits pocilloporids over acroporids (figure 4). In New Caledonia and globally, acroporids contribute exceptionally to coral reef structural complexity, biodiversity, and calcification [13,36,58–60]. Despite high capabilities to dominate reefs in peri-optimal environments, acroporids are particularly sensitive to environmental stressors such as warming and declining water quality, with community shifts from acroporid dominance to pocilloporids or poritids often observed in sub-optimal conditions [7,10,11,29,41]. Widespread acroporid declines have been associated with reef environment degradation in various regions including the Caribbean, Persian Gulf, Great Barrier Reef, and French Polynesia [12,15,16,61]. In contrast to other regions where acroporids appear to face their upper temperature limits [11,29], our study suggests acroporid performances may actually increase in a warmer climate in the sub-tropical reef system of New Caledonia where few large-scale bleaching events have been recorded. Nevertheless, restraining anthropization of coastal environments appears key to preserving near-shore acroporid populations and their unique contributions to reef accretion and resilience.

**Figure 4.**
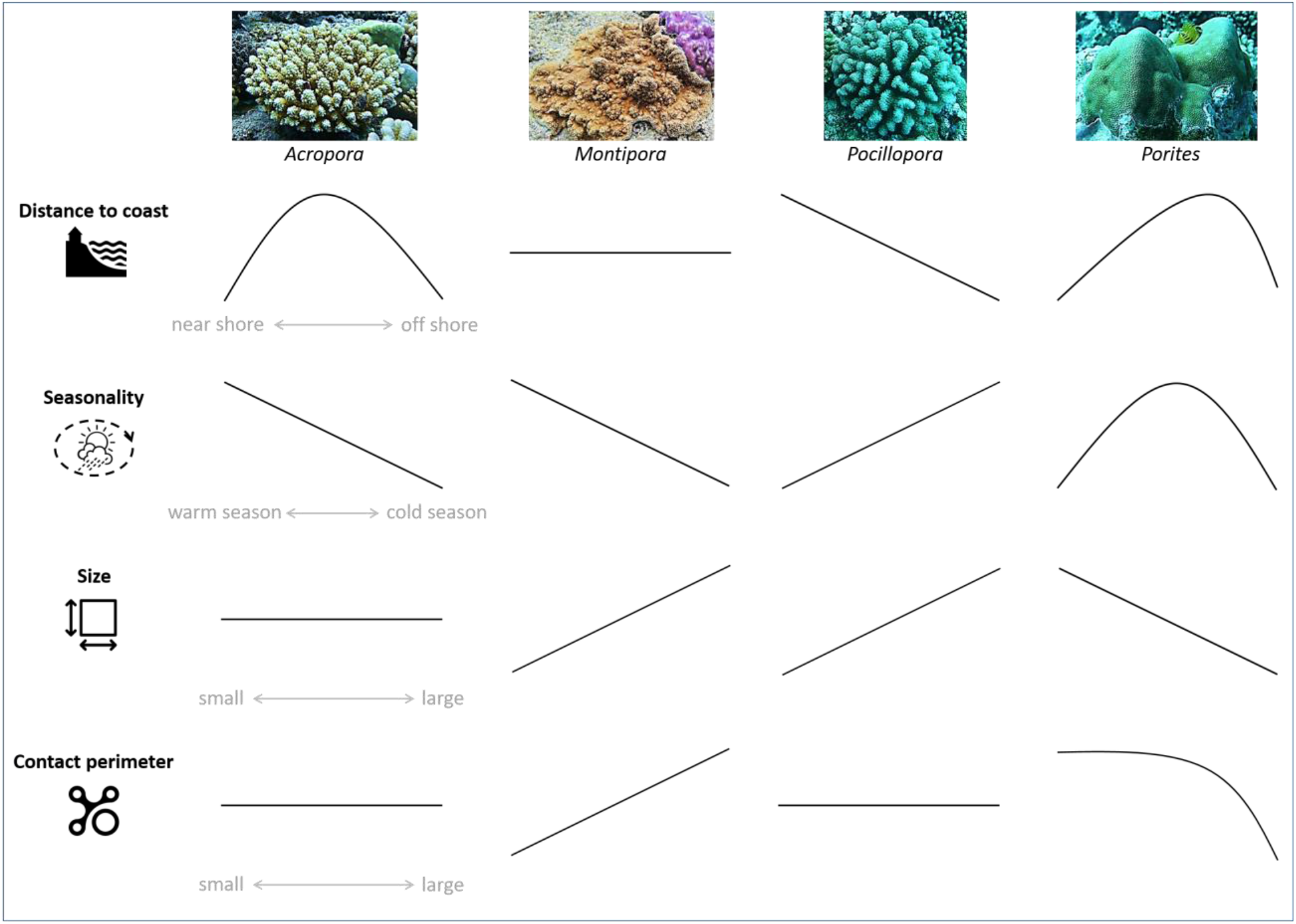
Multi-dimensional niche segregation among the four major reef-building coral taxa as revealed by variability in their competitive performances. The response patterns (summarized from figures 2 and 3) indicate segregation in time and space (different environmental preferences) as well as in life-strategies (different optimal sizes and warfare capacities).

## Conclusions

As global changes increasingly alter coastal marine environments, some ecosystems are inexorably expected to collapse while others may transform to new community compositions, structures, and functions, changes that remain hard to predict [1,3,4,11,]. Nevertheless, present trajectories of coastal degradation indicate future reef environments may increasingly resemble those found near dense human concentrations today [6–8]. Our study suggests that such anthropization results in lower abilities of some major coral taxa in preempting reef space via direct competition. Because corals are slow growing habitat-forming species at the basis of reef ecosystems, such differences in competitive performances may result in extirpations of vulnerable populations, with implications for reef ecosystem biodiversity and services to society.

Overall, competitive performance appears as an effective, widespread, and accessible indicator of species performances across ecological gradients. It can help identify the biological and environmental constraints underlying ecological niches and environmental windows that define species distributions, coexistence, and therefore biodiversity patterns. Given the many ecological pathways that link species performances to their environment, we encourage similar quantitative investigations to further understanding of determinants of species interactions at the interplay between evolutionary traits, life-strategies, and global changes, and implications for the dynamics of ecosystems in a changing environment.

## Acknowledgements

This study was supported by funding from the Laboratoire d’Excellence CORAIL (www.labex-corail.fr) in the form of a 1-year post-doctoral fellowship attributed to Mohsen Kayal. We thank IRD colleagues, Julien Neuveut, and William Roman for assistance for boating and diving. We are grateful to Jane Ballard for major improvements to the manuscript.

