## Supplementary material for "The war of corals: patterns, drivers, and implications of changing coral competitive performances across reef environments"

**Appendix 1.** Further descriptions of coral competitive interactions:

- The war of corals: https://www.instagram.com/p/CFSx-DfBqui/
- When two corals come into contact: https://www.instagram.com/p/CHHJ7X1hCMJ/
- Coral competition at the youngest age: https://www.instagram.com/p/CIwhH7DhF8l/
- Competition in larger corals: https://www.instagram.com/p/CIzfVoKBJw2/
- A battle of acroporids: https://www.instagram.com/p/CKDWsngBHnZ/

**Figure S1.** Distributions of the 20 reefs in the south-western reef system of New Caledonia where coral competitive interactions were inspected. The coastal system is characterized by diverse ecological habitats (fringing, mid-shelf, barrier, and outer-slope reefs) distributed along pronounced cross-shelf environmental gradients in relation to the positions to the coastal city of Noumea and the open ocean. Photograph of the coastal city Noumea by © Mohsen Kayal, satellite image from Google, reef distances to Noumea calculated in R statistical software complemented by package 'geosphere'.

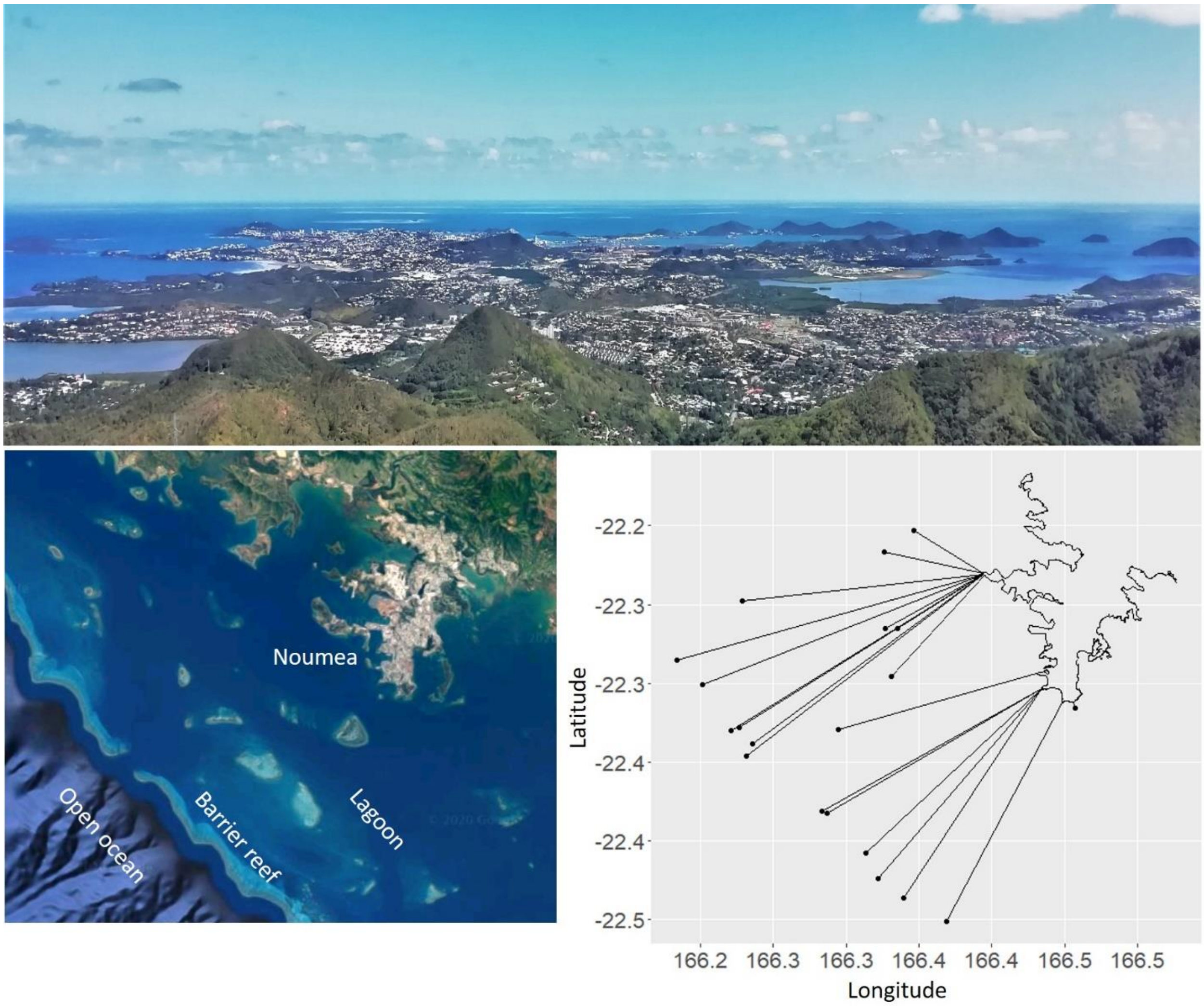

**Figure S2.** Visualization of raw data for some of the variables used in our study. The abundance of focal coral taxa and their competitors as encountered during our surveys are shown (first row), along with their distributions in size (second row) and space (third row), as well as variability in outcomes in terms of overreach and overgrowth (fourth row). Coral competitive interactions were characterized on 20 sites throughout the southwestern reef system of New Caledonia. See table S1 for more information. CCA for crustose coralline algae, BCA for branching coralline algae.

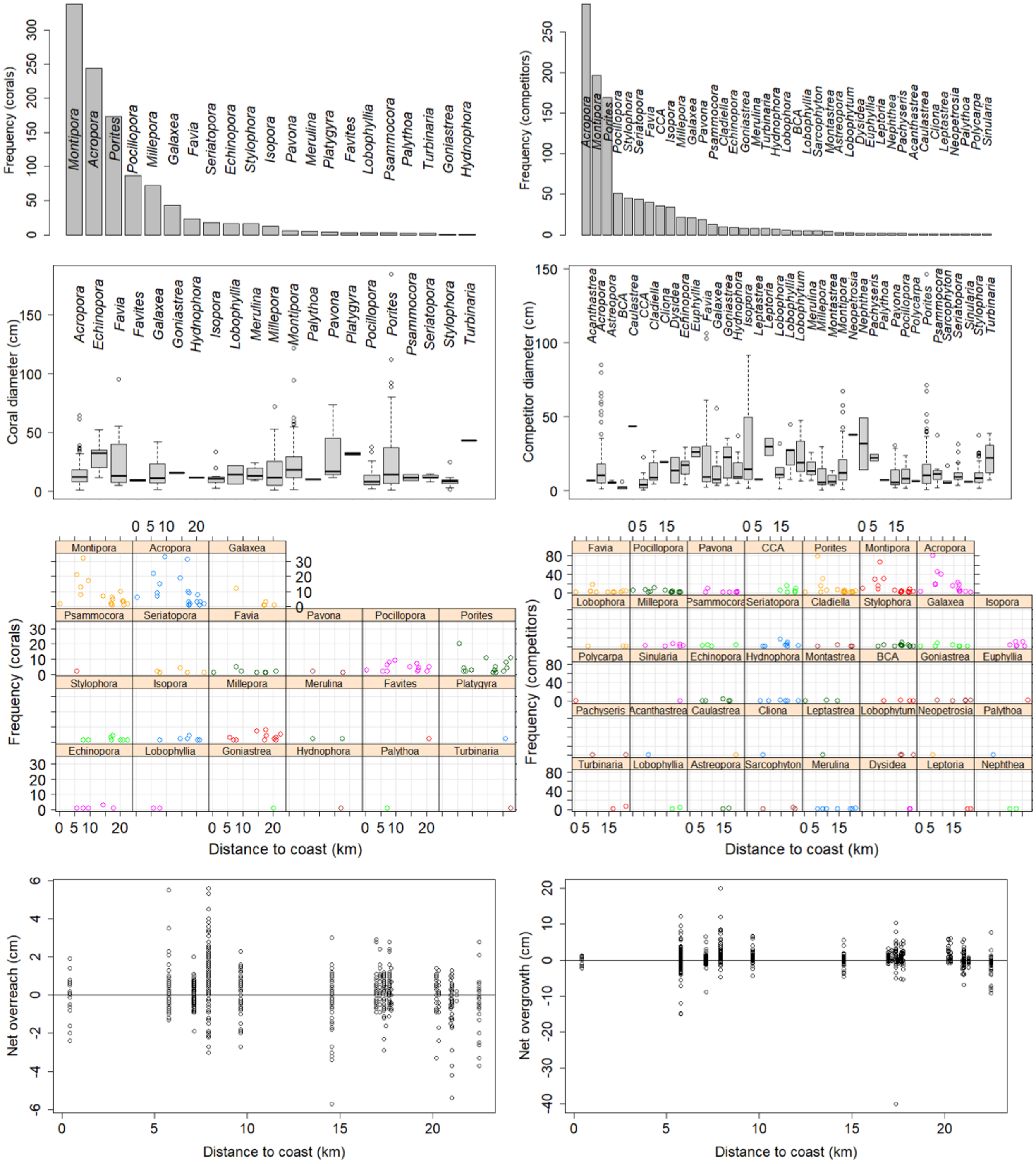

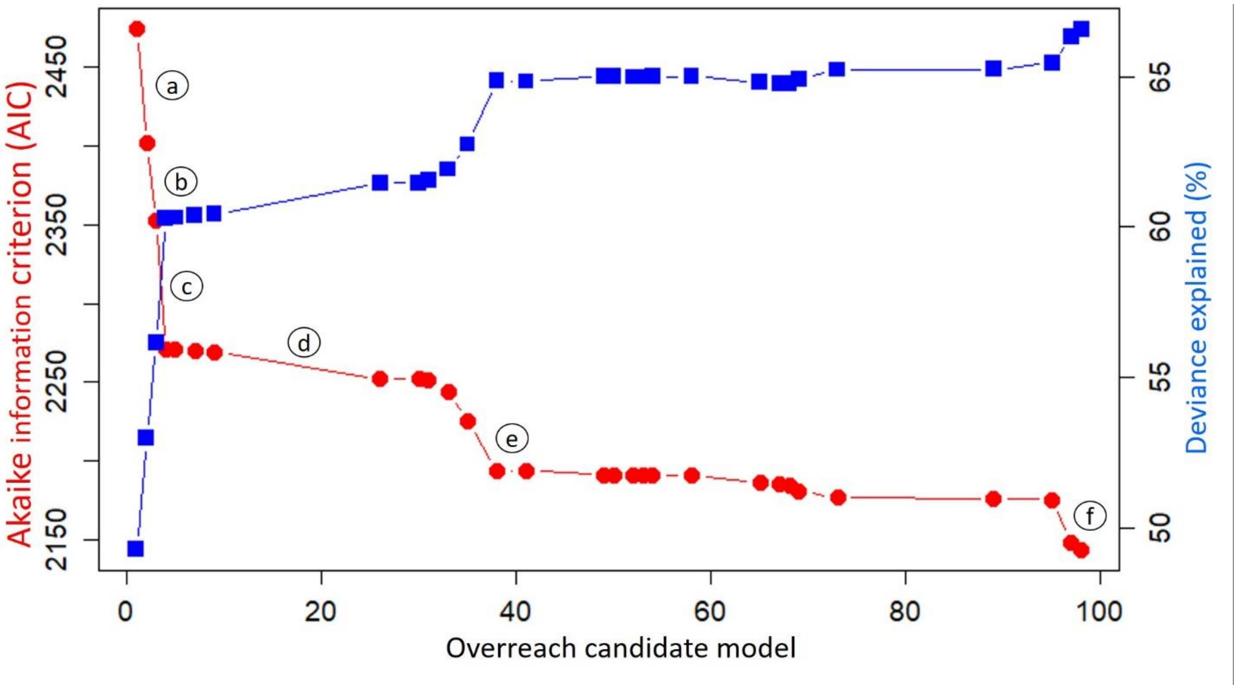


**Figure S3.** Changes in Akaike information criterion and proportion of deviance explained throughout the selection process of candidate generalized additive models (103 total) to characterize coral competitive performances in terms of net overreach. Only significant model selection steps are illustrated. Key steps in the selection process are highlighted: (a) different responses of coral taxa in space, (b) different responses of coral taxa with size, (c) different effects of competitor taxa with competitor size, (d) different responses of coral taxa in time, (e) log-transform over-dispersed covariables, (f) different responses of coral taxa with coral to competitor size-ratio. See table S2 for final model details.

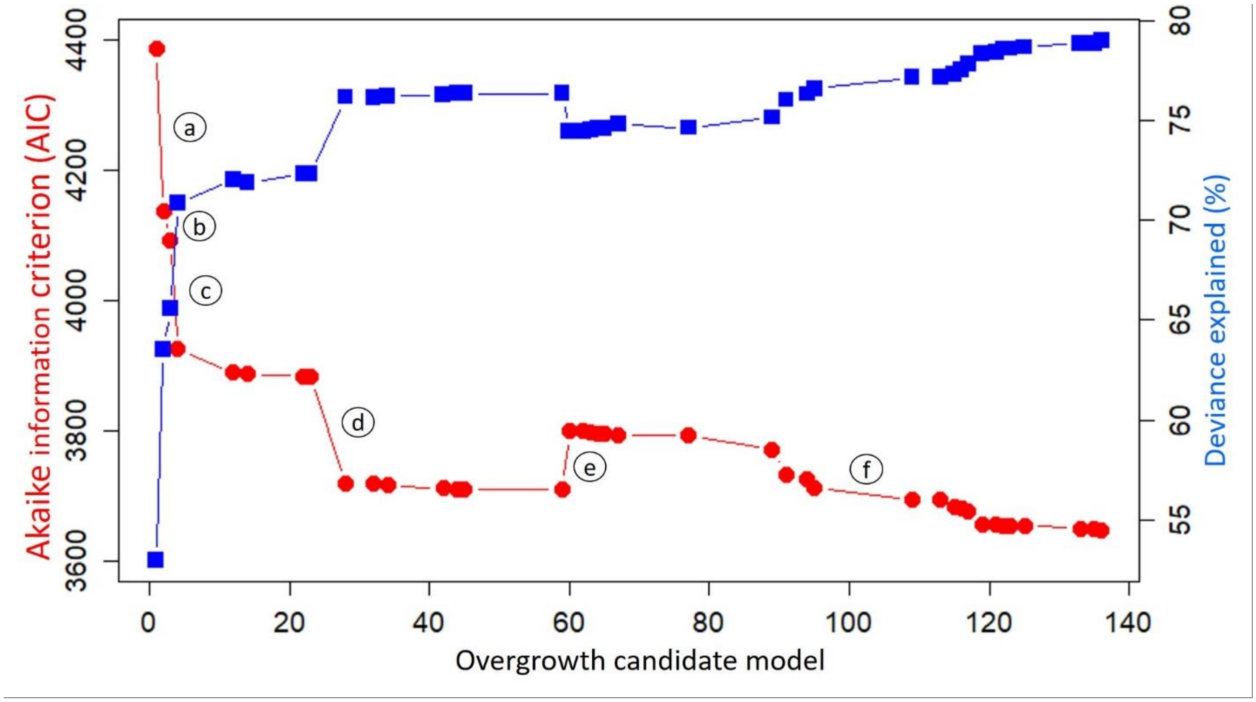


**Figure S4.** Changes in Akaike information criterion and proportion of deviance explained throughout the selection process of candidate generalized additive models (144 total) to characterize coral competitive performances in terms of net overreach. Only significant model selection steps are illustrated. Key steps in the selection process are highlighted: (a) different responses of coral taxa in space, (b) different responses of coral taxa with size, (c) different effects of competitor taxa with competitor size, (d) different responses of coral taxa with contact-perimeter, (e) log-transform over-dispersed covariables, (f) different effects of contact-perimeter in space. See table S3 for final model details.

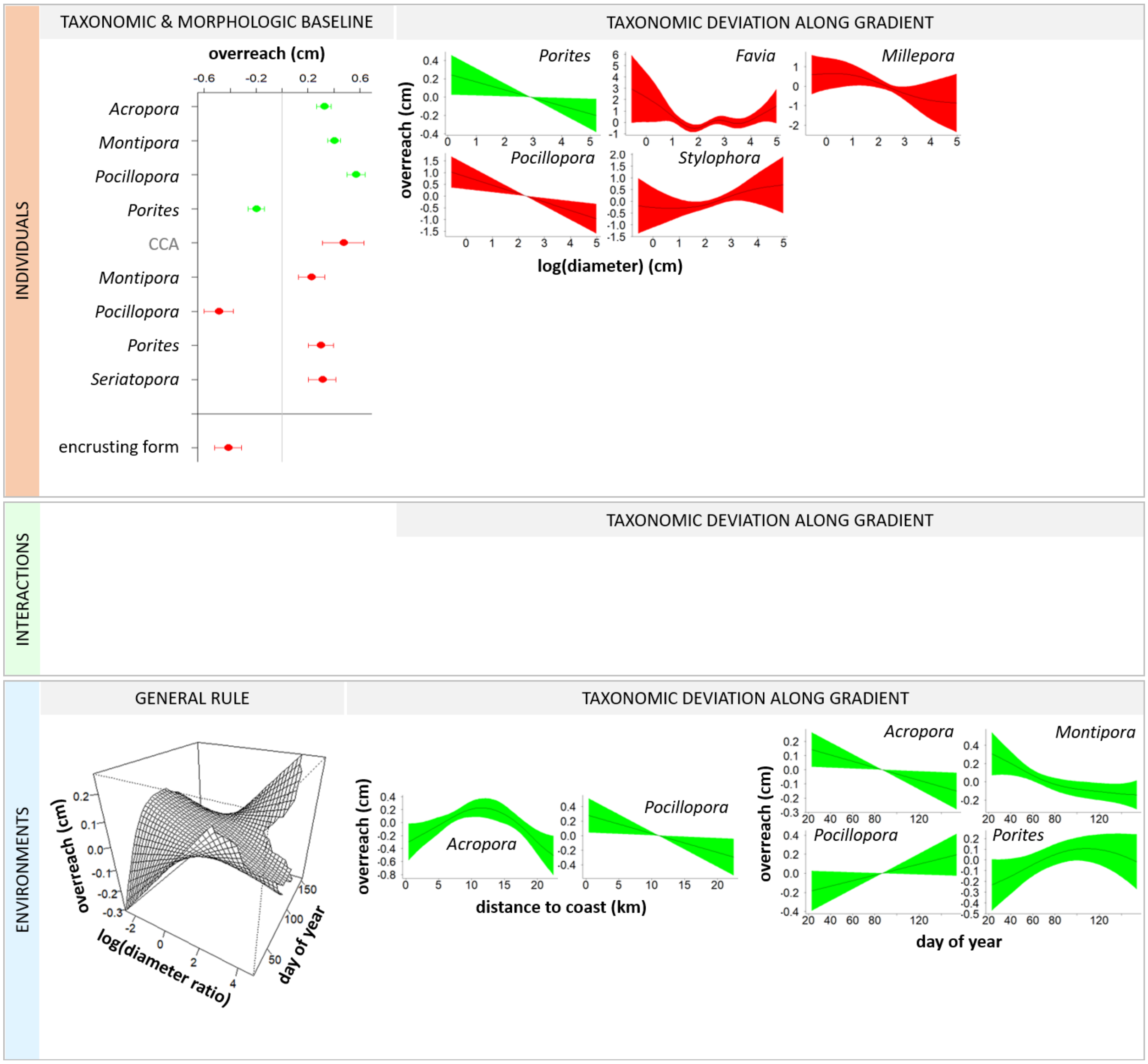


**Figure S5.** Outputs of generalized additive model describing variability in coral net overreach when restricting data to the most abundant taxa. The 5 most abundant coral taxa (*Acropora*, *Montipora*, *Porites*, *Pocillopora*, *Millepora*) and 10 most abundant competitors (*Acropora*, *Montipora*, *Porites*, *Pocillopora*, *Millepora*, *Stylophora*, *Seriatopora*, *Favia*, CCA, *Isopora*) were considered. Similar patterns to those found with the complete dataset (figure 2) confirm the pervasiveness of the mechanisms influencing coral overreach. Only significant effects are illustrated. See figure 2 for details.

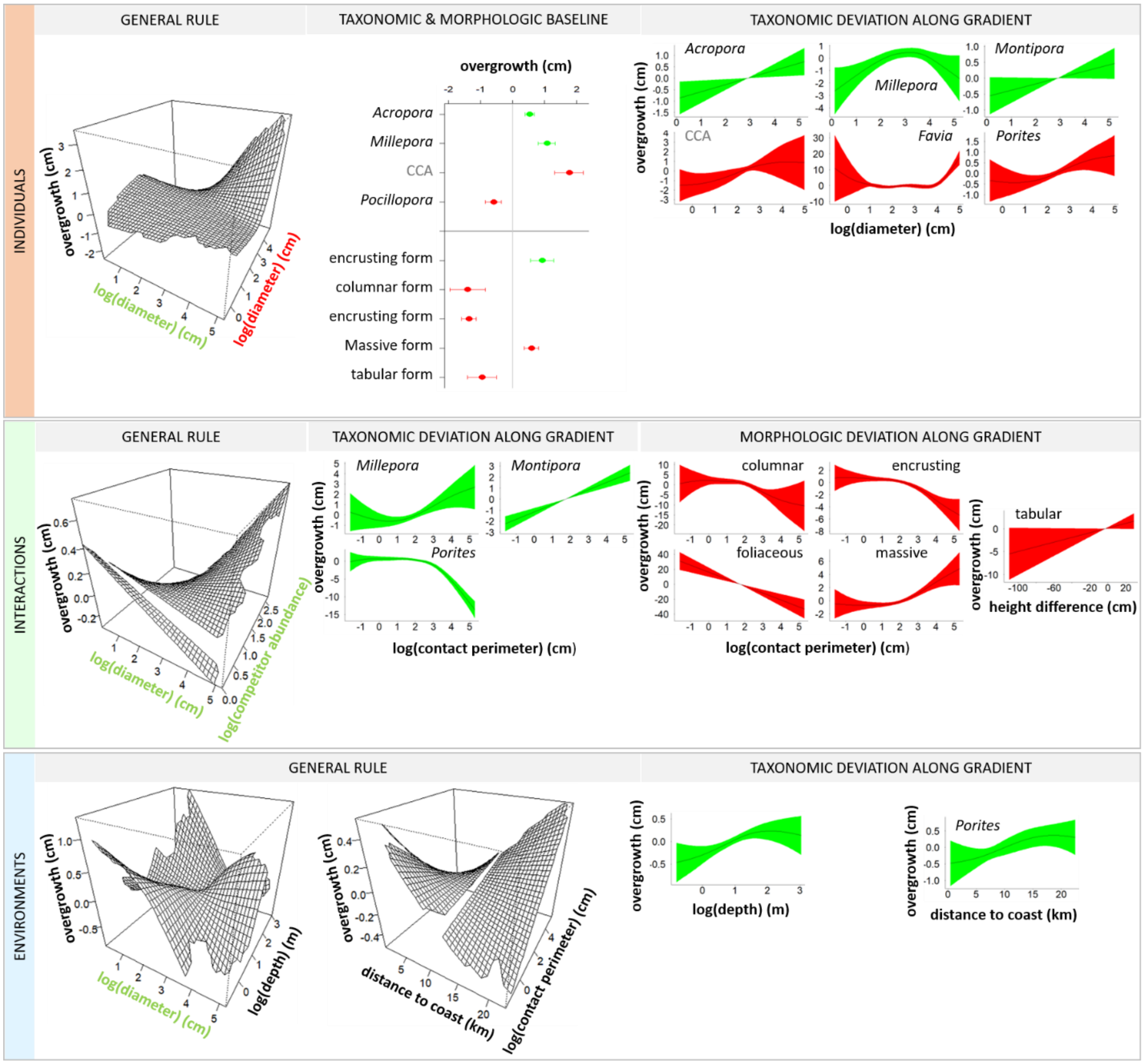


**Figure S6.** Outputs of generalized additive model describing variability in coral net overgrowth when restricting data to the most abundant taxa. The 5 most abundant coral taxa (*Acropora*, *Montipora*, *Porites*, *Pocillopora*, *Millepora*) and 10 most abundant competitors (*Acropora*, *Montipora*, *Porites*, *Pocillopora*, *Millepora*, *Stylophora*, *Seriatopora*, *Favia*, CCA, *Isopora*) were considered. Similar patterns to those found with the complete dataset (figure 3) confirm the pervasiveness of the mechanisms influencing coral overgrowth. Only significant effects are illustrated. See figure 3 for details.

**Table S1.** Variables used to characterize coral competitive performances in terms of net overreach and overgrowth. *In situ* measurements were performed on randomly chosen focal corals from dominant taxa and their surrounding competitor species (see figure S2 for some visualizations of the raw data). Note that the calcifying hydrozoan *Millepora* (fire-coral) was considered among reef-building corals.

| Variable | Encountered levels or range |
|---|---|
| *coral-taxon*<br>(coral taxon at genus level) | Hard corals: *Acropora, Echinopora, Favia, Favites, Galaxea, Goniastrea, Isopora, Lobophyllia, Merulina, Millepora, Montipora, Palythoa, Pavona, Platygyra, Pocillopora, Porites, Psammocora, Seriatopora, Stylophora, Turbinaria, Hydnophora* |
| *comp-taxon*<br>(competitor taxon at different taxonomic levels among groups) | Hard corals: *Acanthastrea, Acropora, Astreopora, Caulastrea, Echinopora, Euphyllia, Favia, Galaxea, Goniastrea, Hydnophora, Isopora, Leptastrea, Leptoria, Lobophyllia, Merulina, Millepora, Montastrea, Montipora, Pachyseris, Pavona, Pocillopora, Porites, Psammocora, Seriatopora, Stylophora, Turbinaria*<br>Soft corals: *Cladiella, Lobophytum, Nephthea, Palythoa, Sarcophyton, Sinularia*<br>Ascidians: *Polycarpa*<br>Sponges: *Cliona, Dysidea, Neopetrosia*<br>Algae: branching coralline algae, crustose coralline algae, *Lobophora* |
| *coral-growth-form* | branching, columnar, encrusting, foliaceous, massive, sub-branching, tabular, vase shape |
| *comp-growth-form* | branching, columnar, encrusting, foliaceous, massive, sub-branching, tabular, vase shape |
| *coral-diameter*<br>(mean diameter in 3 dimensions) | 1.1 – 184.3 cm |
| *comp-diameter*<br>(mean diameter in 3 dimensions) | 0.6 – 146.7 cm |
| *diameter-ratio*<br>(coral / competitor) | 0.07 – 110.6 |
| *contact-perimeter* | 0.2 – 200 cm |
| *height-difference*<br>(competitor - coral) | -110 – 60 cm |
| *n-competitors* | 1 – 15 |
| *total-competing-perimeter* | 0.3 – 283.1 cm |
| *net-overreach* | -5.7 – 5.6 cm |
| *net-overgrowth* | -40 – 20 cm |
| *depth* | 0.45 – 22 m |
| *distance-to-coast* | 0.420 – 22.493 km |
| *day-of-year* | 24/01/2013 – 04/06/2013 |

**Table S2.** Output table of generalized additive model characterizing variability in coral competitive performances in terms of net overreach. s() indicates a smooth term or a smooth-factor interaction, whereas ti() indicates a tensor-product interaction between two smooth terms (Wood 2017). The estimated degrees of freedom (df), F-tests (F), and significance levels (*p*-value) of each model term are shown. Only significant model terms are listed (F>0.1, *p*<0.05).

| **Parameter** | **df** | **F** | ***p*-value** |
|---|---|---|---|
| *coral-taxon* | 20 | 18.19 | < 2e-16 |
| *comp-taxon* | 38 | 8.34 | < 2e-16 |
| *comp-growth-form* | 6 | 5.34 | 1.97e-05 |
| s(log(*coral-diameter*) × *coral-taxon*[*Echinopora*]) | 8.917e-01 | 1.587 | 3.65e-05 |
| s(log(*coral-diameter*) × *coral-taxon*[*Favia*]) | 1.512 | 0.632 | 0.015518 |
| s(log(*coral-diameter*) × *coral-taxon*[*Galaxea*]) | 2.570 | 1.341 | 0.002343 |
| s(log(*coral-diameter*) × *coral-taxon*[*Porites*]) | 1.396 | 0.655 | 0.016697 |
| s(log(*comp-diameter*) × *comp-taxon*[*Favia*]) | 4.645 | 3.435 | 7.29e-06 |
| s(log(*comp-diameter*) × *comp-taxon*[*Galaxea*]) | 4.526 | 5.752 | 7.08e-10 |
| s(log(*comp-diameter*) × *comp-taxon*[*Goniastrea*]) | 3.137 | 4.398 | 5.39e-07 |
| s(log(*comp-diameter*) × *comp-taxon*[*Lobophyllia*]) | 2.516 | 9.837 | 1.95e-09 |
| s(log(*comp-diameter*) × *comp-taxon*[*Merulina*]) | 9.532e-01 | 3.258 | 5.98e-06 |
| s(log(*comp-diameter*) × *comp-taxon*[*Millepora*]) | 1.254 | 0.494 | 0.040386 |
| s(log(*comp-diameter*) × *comp-taxon*[*Montastrea*]) | 9.189e-01 | 3.675 | 0.000538 |
| s(log(*comp-diameter*) × *comp-taxon*[*Pocillopora*]) | 8.685e-01 | 0.784 | 0.007077 |
| s(log(*comp-diameter*) × *comp-taxon*[*Stylophora*]) | 1.923 | 1.268 | 0.002384 |
| s(log(*comp-diameter*) × *comp-taxon*[*Turbinaria*]) | 7.970e-01 | 0.597 | 0.021935 |
| s(log(*comp-diameter*) × *comp-taxon*[*Cladiella*]) | 7.347e-01 | 0.390 | 0.049067 |
| s(log(*comp-diameter*) × *comp-taxon*[*Sarcophyton*]) | 8.136e-01 | 1.111 | 0.018653 |
| s(log(*n-competitors*) × *coral-taxon*[*Echinopora*]) | 9.974e-01 | 7.646 | 9.39e-06 |
| s(log(*n-competitors*) × *coral-taxon*[*Favia*]) | 2.332 | 23.434 | < 2e-16 |
| s(log(*n-competitors*) × *coral-taxon*[*Galaxea*]) | 1.801 | 1.197 | 0.046274 |
| s(log(*n-competitors*) × *coral-taxon*[*Montipora*]) | 2.506 | 0.815 | 0.044263 |
| s(*distance-to-coast* × *coral-taxon*[*Acropora*]) | 2.236 | 1.349 | 0.002896 |
| s(*distance-to-coast* × *coral-taxon*[*Echinopora*]) | 3.217 | 11.911 | 1.12e-11 |
| s(*distance-to-coast* × *coral-taxon*[*Favia*]) | 2.569 | 4.769 | 5.96e-08 |
| s(*distance-to-coast* × *coral-taxon*[*Pocillopora*]) | 1.308 | 1.683 | 0.000147 |
| ti(log(*diameter-ratio*) × *day-of-year*) | 3.214 | 0.628 | 0.004490 |
| s(*day-of-year* × *coral-taxon*[*Acropora*]) | 8.913e-01 | 0.963 | 0.002674 |
| s(*day-of-year* × *coral-taxon*[*Galaxea*]) | 2.739 | 4.222 | 6.16e-07 |
| s(*day-of-year* × *coral-taxon*[*Isopora*]) | 2.990 | 2.717 | 0.000514 |
| s(*day-of-year* × *coral-taxon*[*Montipora*]) | 7.737e-01 | 0.410 | 0.037169 |
| s(*day-of-year* × *coral-taxon*[*Pocillopora*]) | 8.815e-01 | 0.880 | 0.004369 |
| s(*day-of-year* × *coral-taxon*[*Porites*]) | 1.447 | 0.570 | 0.035073 |

**Table S3.** Output table of generalized additive model characterizing variability in coral competitive performances in terms of net overgrowth. s() indicates a smooth term or a smooth-factor interaction, whereas ti() indicates a tensor-product interaction between two smooth terms (Wood 2017). The estimated degrees of freedom (df), F-tests (F), and significance levels (*p*-value) of each model term are shown. Only significant model terms are listed (F>0.1, *p*<0.05). CCA for crustose coralline algae.

| **Parameter** | **df** | **F** | ***p*-value** |
|---|---|---|---|
| *coral-taxon* | 21 | 3.281 | 1.00e-06 |
| *coral-growth-form* | 7 | 3.317 | 0.00171 |
| *comp-taxon* | 37 | 4.733 | < 2e-16 |
| *Comp-growth-form* | 6 | 9.205 | 8.23e-10 |
| ti(log(*coral-diameter*) × log(*comp-diameter*)) | 3.175 | 1.822 | 2.24e-07 |
| s(log(*coral-diameter*) × *coral-taxon*[*Montipora*]) | 7.908e-01 | 0.456 | 0.025134 |
| s(log(*comp-diameter*) × *comp-taxon*[*Echinopora*]) | 2.731 | 1.774 | 0.002872 |
| s(log(*comp-diameter*) × *comp-taxon*[*Favia*]) | 5.342 | 8.977 | 2.87e-15 |
| s(log(*comp-diameter*) × *comp-taxon*[*Goniastrea*]) | 1.474 | 3.160 | 1.15e-06 |
| s(log(*comp-diameter*) × *comp-taxon*[*Hydnophora*]) | 8.373e-01 | 0.978 | 0.015032 |
| s(log(*comp-diameter*) × *comp-taxon*[*Merulina*]) | 7.197e-01 | 0.455 | 0.045375 |
| s(log(*comp-diameter*) × *comp-taxon*[*Porites*]) | 1.744 | 1.501 | 0.000433 |
| s(log(*comp-diameter*) × *comp-taxon*[*Psammocora*]) | 1.944 | 0.944 | 0.021689 |
| s(log(*comp-diameter*) × *comp-taxon*[*Nephthea*]) | 9.968e-01 | 293.971 | < 2e-16 |
| s(log(*comp-diameter*) × *comp-taxon*[CCA]) | 1.094 | 0.506 | 0.027289 |
| s(log(*comp-diameter*) × *comp-taxon*[*Lobophora*]) | 8.873e-01 | 1.456 | 0.003348 |
| ti(log(*coral-diameter*) × log(*n-competitors*)) | 6.500 | 1.670 | 2.30e-05 |
| s(log(*contact-perimeter*) × *coral-taxon*[*Echinopora*]) | 8.180e-01 | 0.545 | 0.017117 |
| s(log(*contact-perimeter*) × *coral-taxon*[*Lobophyllia*]) | 8.811e-01 | 3.756 | 0.003365 |
| s(log(*contact-perimeter*) × *coral-taxon*[*Millepora*]) | 8.684e-01 | 0.777 | 0.005659 |
| s(log(*contact-perimeter*) × *coral-taxon*[*Montipora*]) | 2.052 | 2.129 | 1.32e-05 |
| s(log(*contact-perimeter*) × *coral-taxon*[ *Pavona*]) | 2.368 | 15.543 | 2.94e-16 |
| s(log(*contact-perimeter*) × *coral-taxon*[ *Porites*]) | 4.483 | 22.060 | < 2e-16 |
| s(log(*contact-perimeter*) × *comp-growth-form*[branching]) | 2.076 | 1.845 | 5.68e-05 |
| s(log(*contact-perimeter*) × *comp-growth-form*[columnar]) | 8.174e-01 | 0.625 | 0.018078 |
| s(log(*contact-perimeter*) × *comp-growth-form*[encrusting]) | 1.886 | 2.002 | 2.25e-05 |
| s(log(*contact-perimeter*) × *comp-growth-form*[foliaceous]) | 4.208 | 3.136 | 1.52e-05 |
| s(log(*contact-perimeter*) × *comp-growth-form*[massive]) | 3.694 | 10.155 | < 2e-16 |
| s(*height-difference* × *comp-growth-form*[foliaceous]) | 2.194 | 1.269 | 0.003043 |
| s(*height-difference* × *comp-growth-form*[ massive]) | 9.509e-01 | 0.388 | 0.045852 |
| ti(log(*coral-diameter*) × log(*depth*)) | 4.176 | 0.818 | 0.002267 |
| ti(*distance-to-coast* × log(*contact-perimeter*)) | 6.311 | 2.851 | 2.51e-09 |
| s(*distance-to-coast* × *coral-taxon*[*Echinopora*]) | 1.968 | 3.095 | 0.000741 |
| s(*distance-to-coast* × *coral-taxon*[*Porites*]) | 2.094 | 2.012 | 9.76e-05 |
